# Evaluating 6- and 18-hour stimulation durations for natural killer cell degranulation (CD107a assay) to optimize workflow efficiency in a clinical immunology laboratory

**DOI:** 10.64898/2026.03.02.708872

**Authors:** Lyndsey Feehan, Leslie Koutoufaris, Jeri Dorsey, Michele Paessler, Priyanka Pandey

**Affiliations:** Clinical Immunology Laboratory, Abramson Research Center, Children’s Hospital of Philadelphia, Philadelphia; Department of Pathology and Laboratory Medicine, Perelman School of Medicine, University of Pennsylvania

## Abstract

**Background:** Natural killer (NK) cell degranulation is a key immune defense mechanism where exposure to tumor or virus-infected cells triggers the fusion of cytoplasmic granules containing apoptotic proteins, perforin, and granzyme with the cell membrane. This process transiently expresses CD107a on the NK cell surface, and measuring CD107a is a standard method to assess NK cell activity.

**Methods:** We compared two stimulation protocols differing only in duration (6-hour vs. 18-hour) using K562 target cells to induce NK cell degranulation. Isolated PBMCs without stimulation served as controls to assess spontaneous degranulation. Anti-CD107a-PE antibody was present throughout stimulation in both test and control samples. After stimulation, cells were stained with anti-CD45, anti-CD3, and anti-CD56 and analyzed by flow cytometry.

**Results:** For 6 of 7 healthy controls, results from both methods fell within 2 standard deviations. Notably, longer (18-hour) stimulation resulted in lower CD107a expression than the 6-hour assay. Interlaboratory comparisons of two samples showed no significant difference (p>0.05). In a suspected hemophagocytic lymphohistiocytosis (HLH) case, two labs reported similarly reduced CD107a expression (9% and 7%). Inter-day variability was observed in a donor across both time points. The 6-hour assay showed higher sensitivity and specificity than the 18-hour assay. A resting period before ex vivo PBMC assays was found necessary.

**Conclusion:** Stimulation periods beyond 6 hours are unsuitable for clinical NK degranulation assays. Screening for HLH should include multiple stimulants to improve assay reliability.

## Introduction

Lysosomal-associated membrane protein-1 (LAMP-1), commonly known as CD107a is marker for effective natural killer (NK) cell degranulation [1–4]. Exposure to foreign antigens lead to the activation of NK cells causing the release of pore-forming protein, perforin and apoptosis causing serine protease, granzyme B [5]. Together, these two proteins act to kill the target cells expressing non-self or foreign antigens on their surfaces. In the process, CD107a protein which colocalizes with perforin and granzyme in the lysosomal granules of NK cells is expressed on the cell surface [6].

The degranulation mechanism has been found to be defective in hemophagocytic syndromes such as Familial hemophagocytic Lymphohistiocytosis (HLH) type 3, 4, 5, Griscelli syndrome, Chediak-Higashi Syndrome, and Hermansky-Pudlak syndrome type II. These conditions are characterized by an overactive immune system that leads to excessive inflammation and debilitating illness in affected patients [7, 8]. HLH-2004 diagnostic criteria established low, or no NK cell activity as one of the criteria for the HLH diagnosis [9].

In HLH-2024 criteria, utilization of NK cell function tests in acute settings was deemphasized because the testing has been technically demanding and required specialized laboratory resources which at times causes delays due to limited accessibility [10]. However, in immunology clinics, the NK cell function tests continue to be part of initial HLH workup in all suspected HLH cases, to differentiate primary from secondary HLH, research and phenotyping of immune dysfunction, and long-term immunologic follow-up. While CD107a degranulation assay alone could only provide information about defective degranulation of NK cells, however, in conjunction with NK cell cytotoxicity, perforin-granzyme, and SAP/XIAP assay, it can serve as an effective tool for screening HLH patients prior to further confirmatory testing. It is important to note that, successful degranulation does not entail complete elimination of target cells; it just depicts the effective working of degranulation mechanism [7, 11, 12].

CD107a degranulation assays are typically performed by incubating peripheral blood mononuclear cells (PBMCs) isolated from patients and healthy controls with MHC-deficient K562 cells along with anti-CD107a [3, 13]. The addition of anti-CD107a to the reaction mix during stimulation periods is necessary to capture transiently expressed CD107a on the NK cell surface, as it is internalized rapidly. [14]. Depending on laboratory protocols, protein transport inhibitors such as monensin or brefeldin may also be included [15]. Subsequently, the reaction mix is washed, and surface stained with antibodies to the markers CD45, CD3, and CD56. Flow cytometry is performed to assess CD107a positivity on CD45^+^CD3^−^CD56^+^ NK cells, with a positivity rate of ≤ 5-10% indicating defective or diminished degranulation [3, 11].

Various research groups have investigated different incubation durations for the CD107a assay by stimulating PBMCs or purified NK cells with K562 cells, experimenting with time frames ranging from 2 to 18 hours [6, 16–18]. In our study, we specifically evaluated the efficacy of the CD107a degranulation assay by exposing PBMCs from healthy controls to K562 cells for a standard 6-hour period, which is commonly used in clinical settings, and an extended duration of 18 hours.

Since clinical samples can arrive at any time during the day, in a standard 6-hour NK cell degranulation assay, PBMCs are isolated immediately upon receipt of blood samples, rested overnight at 37□/5%CO_2_, while the incubation with K562 cells takes place on Day 2. With a 6-hour incubation on Day 2, the clinical staff typically requires more than 8 hours to complete the assay, delaying results until Day 3. Conversely, initiating the incubation of K562 cells with isolated PBMCs on Day 1 can expedite the reporting of results to early Day 2, thereby reducing the workload for staff.

Recognizing the variability in sample arrival times at clinical laboratories, our objective was to streamline the workflow and create a more user-friendly assay that meets the operational needs of a clinical immunology laboratory. Therefore, we focused solely on comparing the 18-hour incubation period against the established 6-hour period. This approach ensures that laboratories can efficiently manage their processes while upholding the accuracy and reliability of their assays.

## Materials and Methods

### Ethics statement

The study was approved by The Children’s Hospital of Philadelphia Institutional Review Board.

### Antibodies

The following antibodies were utilized in this study: PacOrange (PO) anti-human CD45-(clone 2D1, Sysmex), PE anti-human CD107a (clone H4A3, BD Biosciences), PE/Cy5.5 anti-human CD3-(clone UCHT1, Beckman Coulter), and APC anti-human CD56 (clone NCAM16.2, BD Biosciences). Prior to their incorporation into the study, antibodies were titrated to determine optimal concentrations for effective staining.

### PBMC isolation

PBMCs were isolated through density gradient centrifugation using Ficoll-Paque™ PLUS (Cytiva). Blood samples were processed within 24 hours post-collection. Briefly, whole blood collected in sodium heparin anticoagulant vacutainer was mixed with equal volume of complete RPMI medium, which consists of RPMI1640 supplemented with 10% fetal bovine serum (FBS, R&D Systems) and 1% penicillin/streptomycin (Gibco). The diluted blood was layered over density gradient media in a Sepmate™ tube (Stemcell technologies) and centrifuged at 1200g for 10 minutes in a swinging bucket centrifuge. The supernatant was collected and washed with media at 300g for 10 minutes. The resulting pellet was re-suspended in complete RPMI medium, counted, and assessed for viability using 7-AAD on NovoCyte Penteon flow cytometer (Agilent Technologies).

### NK cell degranulation assay

#### 6-hour protocol (Figure 1)

1. Day 0 (resting phase): a 5×10^6^ PBMCs/mL were incubated overnight at 37°C/5% CO_2_.
2. Day 1 (stimulation phase): a 5×10^6^ PBMC/ml were co-cultured with K562 cells at 1:1 effector to target ratio along with PE anti-CD107a in a total reaction volume of 220 µL for 6 hours at 37°C/5% CO_2_.

#### 18-hour protocol (Figure 2)

1. Day 0 (stimulation phase): a 5×10^6^ PBMC/ml were co-cultured with K562 cells at 1:1 effector to target ratio along with PE-anti-CD107a in a total reaction volume of 220 µL for 18 hours at 37°C/5% CO_2_.

Following incubation, cells were washed with 1X phosphate buffer saline (PBS) containing 1% FBS and surface stained with PO anti-CD45, PE/Cy5.5 anti-CD3, and APC anti-CD56 for 30 minutes at 4°C. After staining, cells were washed and resuspended in 300µL PBS with1%FBS. Samples were analyzed using a NovoCyte Penteon flow cytometer (Agilent Technologies) under the following acquisition settings: Time = 3 minutes, Volume = 250µL, CD56+ NK cell gate = 4000 events, and Threshold = 250,000 on FSC-H.

Cell count and viability assessments were performed on both Day 0 and Day 1 for each protocol. All incubations were conducted in 12 × 75 mm polypropylene tubes with loosened caps to facilitate gas exchange. Tubes were placed on slant racks during overnight incubations to maximize the surface area available to cells.

### Inter-laboratory comparison

Parallel testing was performed on two whole blood samples collected in sodium heparin vacutainers. The whole blood samples were shipped at room temperature via overnight delivery to the Diagnostic Immunology Laboratories at CCHMC, in accordance with specimen acceptability guidelines. The lab at CCHMC performs NK cell degranulation assays as part of routine clinical testing (https://www.testmenu.com/cincinnatichildrens/Tests/660971) and utilizes 6-hour incubation protocol. In the present study, the 6-hour protocol was adopted from the said laboratory. While the samples were being shipped, our lot of samples remained at room temperature on the bench top. Both laboratories conducted the assays on the same day.

### Patient sample

A suspected central nervous system HLH case (10-year-old, male) sample was evaluated for NK cell degranulation both at CCHMC and in our laboratory. Due to time and staffing constraints, we froze the isolated PBMCs from the patient’s blood and performed CD107a degranulation assay the following week. The patient sample was tested alongside a healthy control (drawn on the same day as patient), processed, and PBMCs frozen along with patient sample. Due to paucity of the sample, we tested the patient sample and relevant control only with 18-hour protocol, as CCHMC had run the sample using the 6-hour protocol.

### Assay sensitivity and specificity

To identify the minimum amount of antibody necessary to effectively detect CD107a during the degranulation process of NK cells (sensitivity) and that signal produced during the assay was due to the specific binding of PE anti-CD107a (specificity), a competitive environment was created by diluting PE anti-CD107a in unconjugated anti-CD107a at 1:2, 1:4, 1:8, 1:16, and 1:32 dilutions. The diluted antibodies at various concentrations were added to the reaction mix containing PBMC and K562 cells, with evaluations conducted over both 6-hour and 18-hour time periods.

### Data analysis

Data analysis was conducted using FCS Express (De Novo software), Microsoft Excel (single-factor ANOVA), and DATAtab [19].

## Results

### Gating strategy and cursor placement

Lymphocytes were gated based on high CD45 (x-axis) expression and side scatter (SSC-H, y-axis). Doublets were excluded from lymphocyte gate using forward scatter height vs area. Single cells were gated on CD3 versus CD56 to isolate CD3^−^CD56^+^ NK cells. CD107a expression was then evaluated on these NK cells (Figure 3). It was noted that cursor placement for marking CD107a positive expression could be challenging due to the positively skewed distribution in stimulated tubes. Minor adjustments in marker placement could lead to significant shifts in CD107a expression reporting. To enhance data visualization and cursor placement on histograms, density plots were generated with CD107a on the x-axis and side scatter (SSC-H) on the y-axis (Figure 4A and 4B). It was also observed that histograms viewed in an overlaid format presented a different marker placement than those viewed in a stacked format (Figure 4C). For consistency in data analysis, we utilized the subtraction plot function in FCS Express, [20], which automatically calculated the percentage of CD107a-positive NK cells by subtracting the background expression observed in the unstimulated (negative control) tubes from that in the stimulated (test sample) tubes. (Figure 4D).

**Figure 1.**
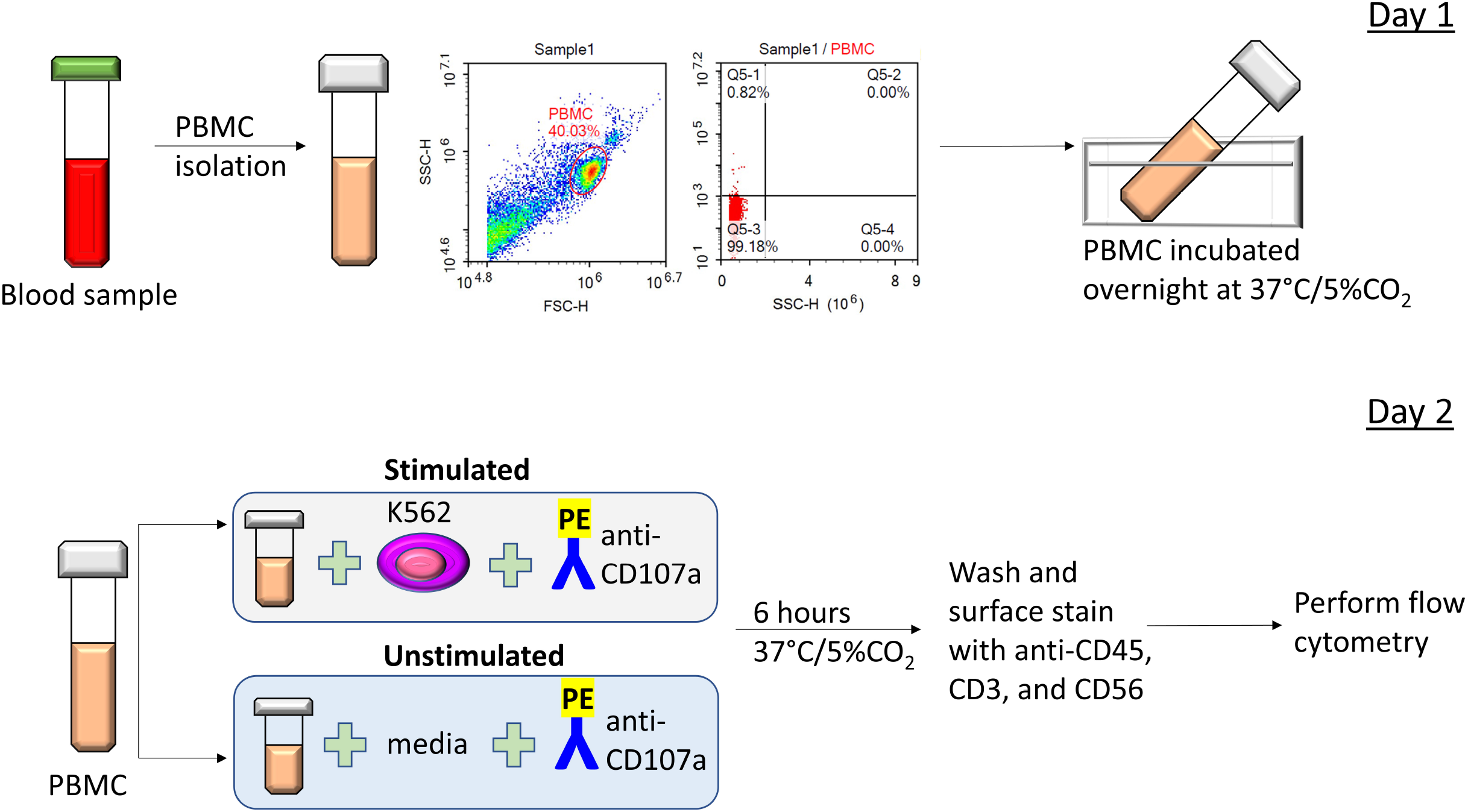
Workflow 6-hour time point. Peripheral blood was collected in sodium heparin tubes and used for the isolation of PBMCs. Following isolation, cells were counted, and viability was assessed using 7-AAD dye. A minimum viability of 85% was required to proceed to Day 2. PBMCs were then resuspended at a concentration of 5×10 cells/mL in complete RPMI medium. The cell suspension was incubated overnight at 37°C with 5% CO in a 12×75 mm polypropylene tube positioned at a 45-degree slant to increase surface area and prevent cell clumping. The following day, PBMCs were washed, counted, and reassessed for viability. Samples with ≥85% viable cells continued to the next step. PBMCs were co-incubated with K562 cells at a 1:1 ratio in the presence of PE-conjugated anti-CD107a antibody for 6 hours at 37°C with 5% CO □. A control tube containing PBMCs and PE-conjugated anti-CD107a (without K562 cells; complete RPMI used instead) was incubated in parallel under identical conditions. After incubation, the reactions were halted by washing the cells. Both test and control samples were then stained with the following surface markers for 30 minutes at 4°C: Pacific Orange anti-CD45, APC anti-CD56, and PE-Cy5.5 anti-CD3. After staining, cells were washed and resuspended in 300 µL cold PBS containing 1% heat-inactivated fetal bovine serum (FBS). Flow cytometry analysis was performed using Agilent’s NovoCyte Penteon. Throughout incubations at 37°C/5% CO□, tube caps were kept slightly ajar to allow for gas exchange. Reactions were stopped upon reaching one of the following criteria: acquisition of 10,000 CD56 events, or a total of 3 minutes, or a volume of 250 µL. The forward scatter height (FSC-H) threshold was set at 75,000. FBS – heat-inactivated fetal bovine serum; PBMC – peripheral blood mononuclear cells; PBS – phosphate-buffered saline.

**Figure 2.**
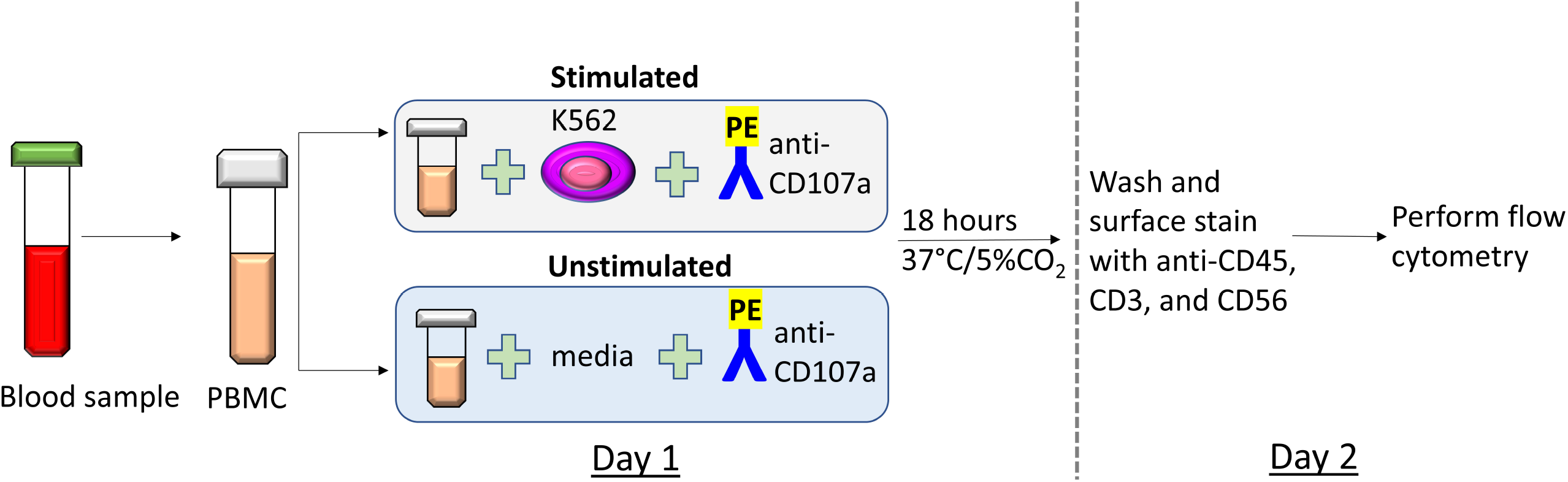
Workflow 18-hour time point. The protocol for isolating PBMCs on Day 1 remains consistent for both the six-hour and eighteen-hour time points. The key distinction between these two time points lies in the fact that the incubation of PBMCs with K562 cells occurs on the same day as their isolation. This approach aims to optimize the workflow by integrating the resting and stimulation phases of the PBMCs. On Day 2, the PBMCs are washed, and flow cytometry is conducted following the procedure outlined in Figure 1.

**Figure 3.**
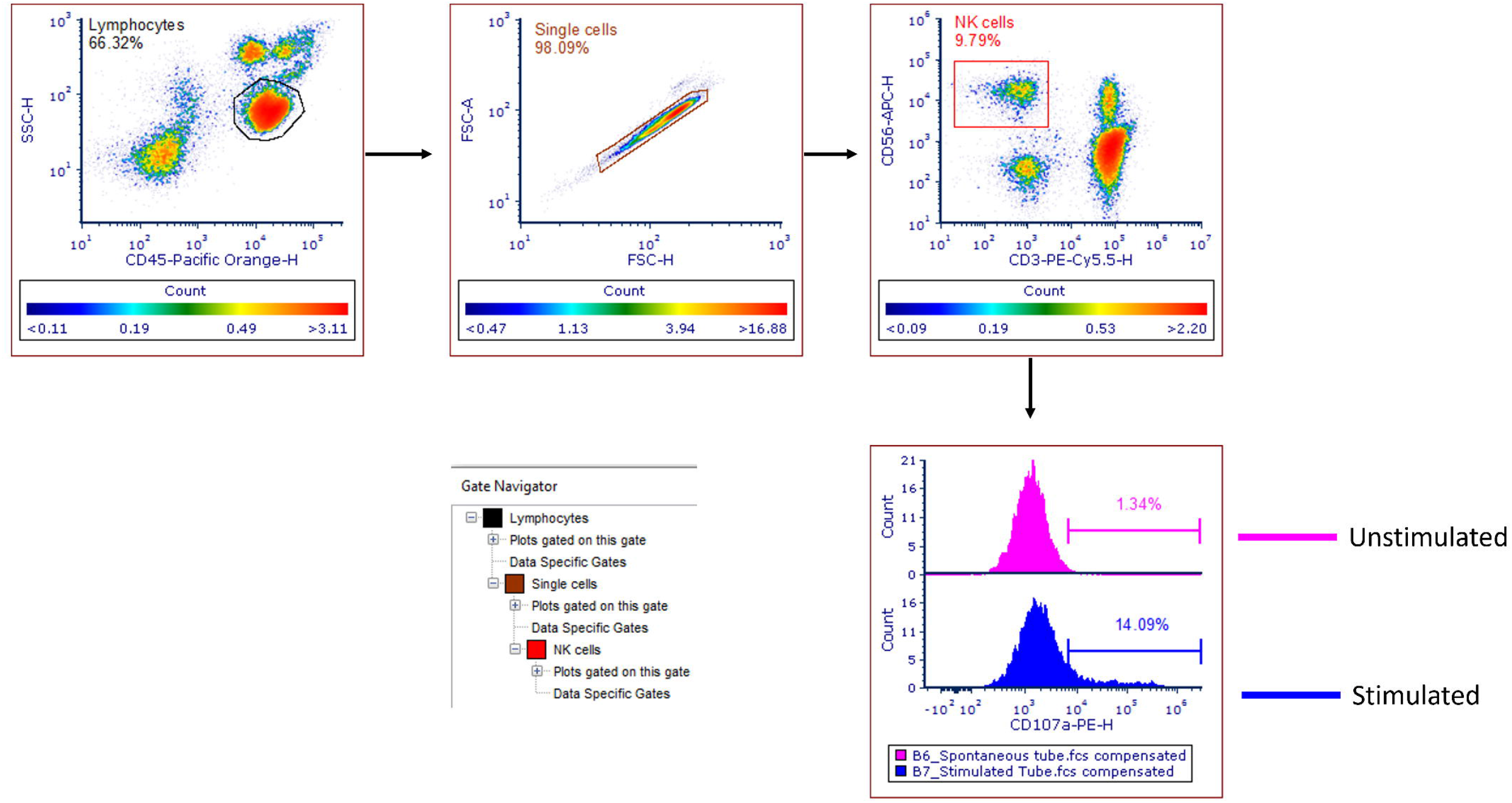
Gating strategy. Lymphocytes were gated on the SSC-H versus CD45 axis. To eliminate doublets from the analysis, lymphocytes were further plotted on the FSC-A versus FSC-H axis. The single cells were then analyzed on the CD56 versus CD3 axis. Subsequently, CD56^high^CD3^negative^ NK cells were identified to assess the expression of CD107 on the surface of NK cells, which serves as a marker for NK cell degranulation. FSC-A: forward scatter-area, FSC-H: forward scatter-height, NK: natural killer, SSC-H: side scatter-height.

**Figure 4.**
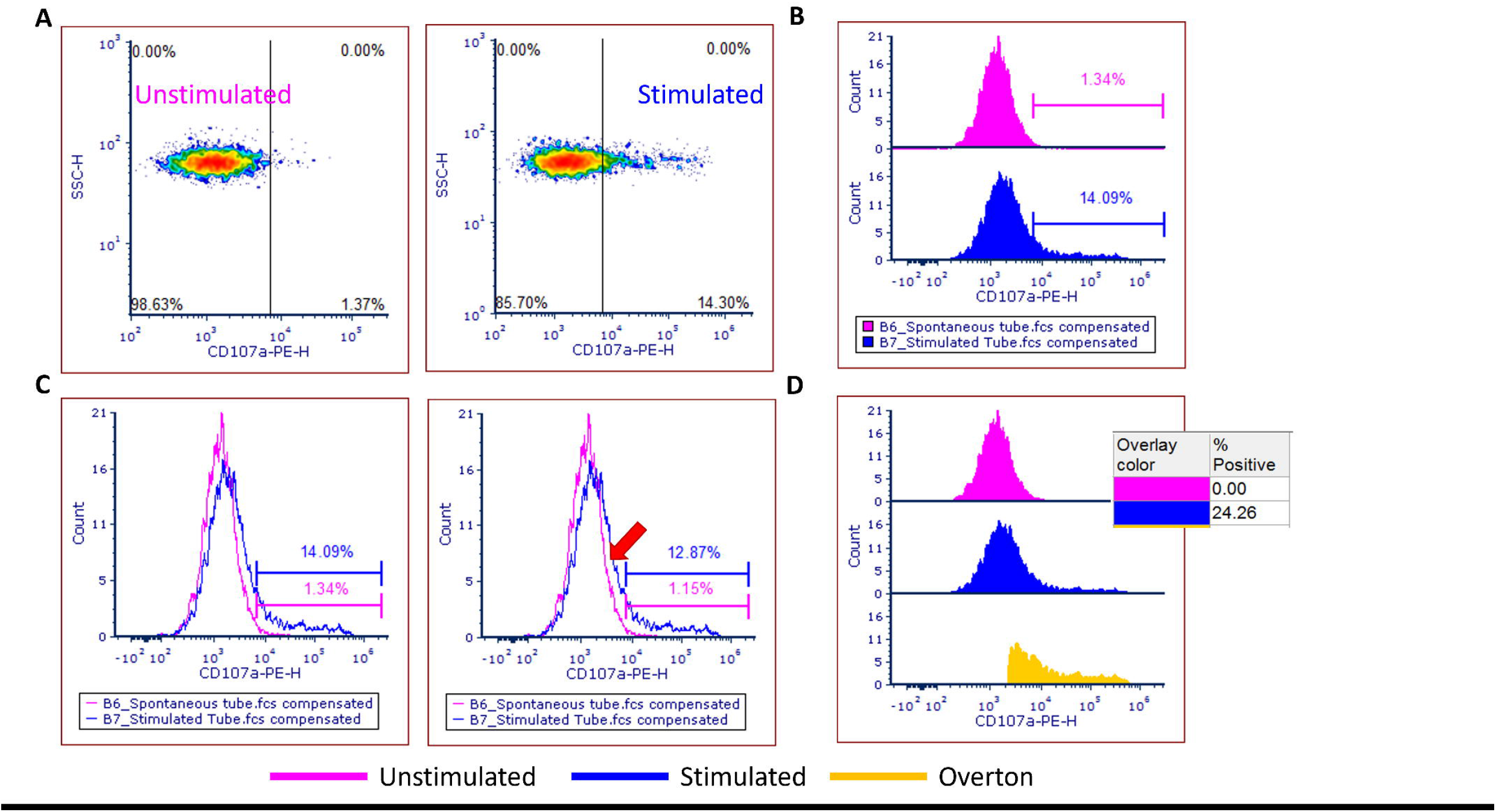
Cursor placement. In assays such as CD107a, accurate cursor placement—specifically, drawing markers to visualize CD107a expression—is critical for reliable result reporting. Markers were initially set on unstimulated samples based on visual identification of the dense population’s endpoint. A. Data was first visualized using density plots. B. The same data was also shown as stacked histograms, with histogram markers aligned using axis references from the density plots. C. When the histograms were overlaid and markers were placed in the same position as on the density plots, D. We realized that, had we relied solely on the histogram view without referencing the density plot, we would have placed the markers differently resulting in a change in CD107a positivity from 14.09% to 12.87%. While this difference may appear minor, it becomes significant given that the cut-offs for CD107a positivity are 11% for healthy subjects and 10% for diseased individuals. E. To address this issue and eliminate manual bias, we automated the analysis by using subtraction plots. In these, CD107a positivity (%) is calculated as the difference between stimulated and unstimulated expression levels. Subtraction plots were generated using FCS Express.

### 6-hour vs 18-hour

A Bland-Altman plot was generated using data from 7 healthy control samples to compare the two protocols. The 6-hour protocol showed higher NK cell degranulation, with CD107a positivity averaging 17±5.6% (mean ± SD), compared to 10±4.5% with the 18-hour protocol. The 18-hour method consistently produced lower CD107a expression, as indicated by all 7 data points falling below the zero-difference line. Except for one point, all values were within ±2 standard deviations—the 95% limits of agreement. However, the range of the 95% confidence interval was notably wide (from –22.19 to +5.33), indicating poor agreement between the two protocols. (Figure 5).

**Figure 5.**
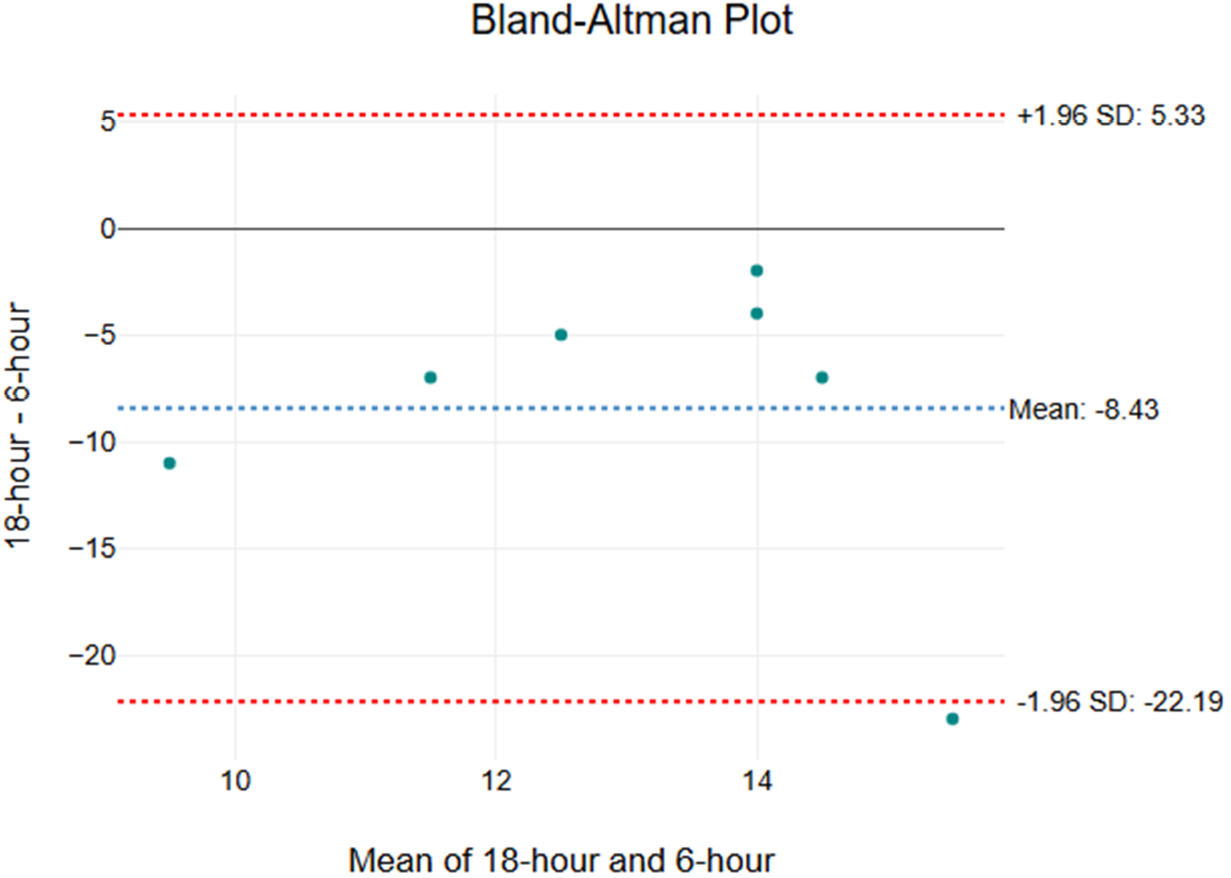
Method comparison. A Bland-Altman plot was created to compare 6- and 18-hour protocols. The mean difference (bias) between the two methods was −8.43, indicating that 18-hour values were consistently lower i.e. 18-hour protocol was consistently generating CD107a positivity or NK cell degranulation less than 6-hour protocol. The limits of agreement range from −22.19 to 5.33, suggesting substantial variability between the methods. Six out of 7 data points were within the limits of agreement (95% CI), 1 data point falling out of the confidence interval was not an extreme outlier.

### Inter-laboratory comparison

There was no significant difference in the CD107a expression between CCHMC 6-hour, CHOP 6-hour, and CHOP 18-hour protocols (p>0.05, n=2), Figure 6.

**Figure 6.**
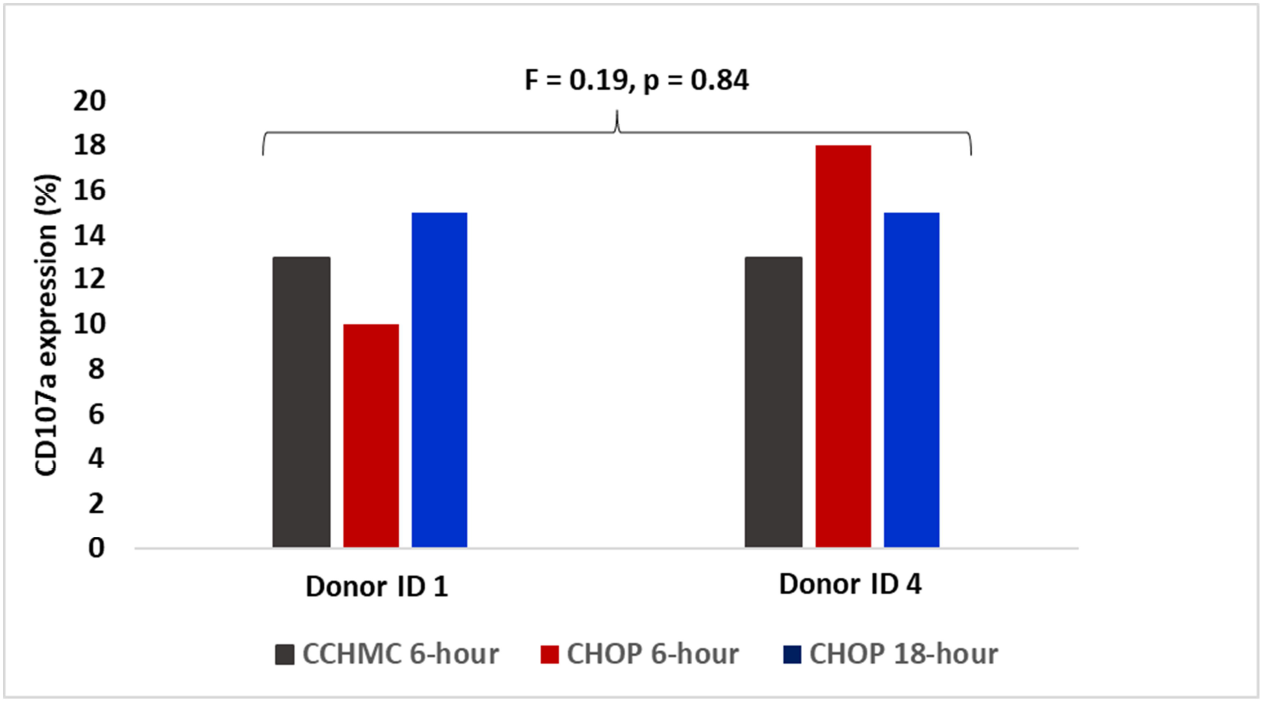
Inter-laboratory comparison. No statistically significant difference was found between the two samples across the three protocols conducted in the two laboratories (one-way ANOVA, p > 0.05). The variation observed between the CHOP-6 hour and CCHMC protocol values may be due to differences in the flow cytometers used for data acquisition. CHOP: The Children’s Hospital of Philadelphia, CCHMC: Cincinnati Children’s Hospital Medical Center.

### Patient sample run

The NK cells from the suspected HLH case exhibited low CD107a expression at 8.77% while the frozen healthy control showed 17.15% CD107a expression, indicating a normal level of NK cell degranulation typically seen in healthy individuals (Figure 7). CCHMC recorded a CD107a expression of 7% for this patient sample. A validated cutoff of 10% for NK cell degranulation has been established for this assay, suggesting that any samples exceeding this threshold are regarded as having normal NK cell activity [3]. The patient demonstrated normal NK cell function (as assessed by chromium release assay) and perforin/granzyme expression. CNS HLH was considered but ruled out as were autoinflammatory causes. The subject was treated with various immune therapies such as steroids, IVIG, Rituximab, and Cyclophosphamide, which helped control his symptoms.

**Figure 7.**
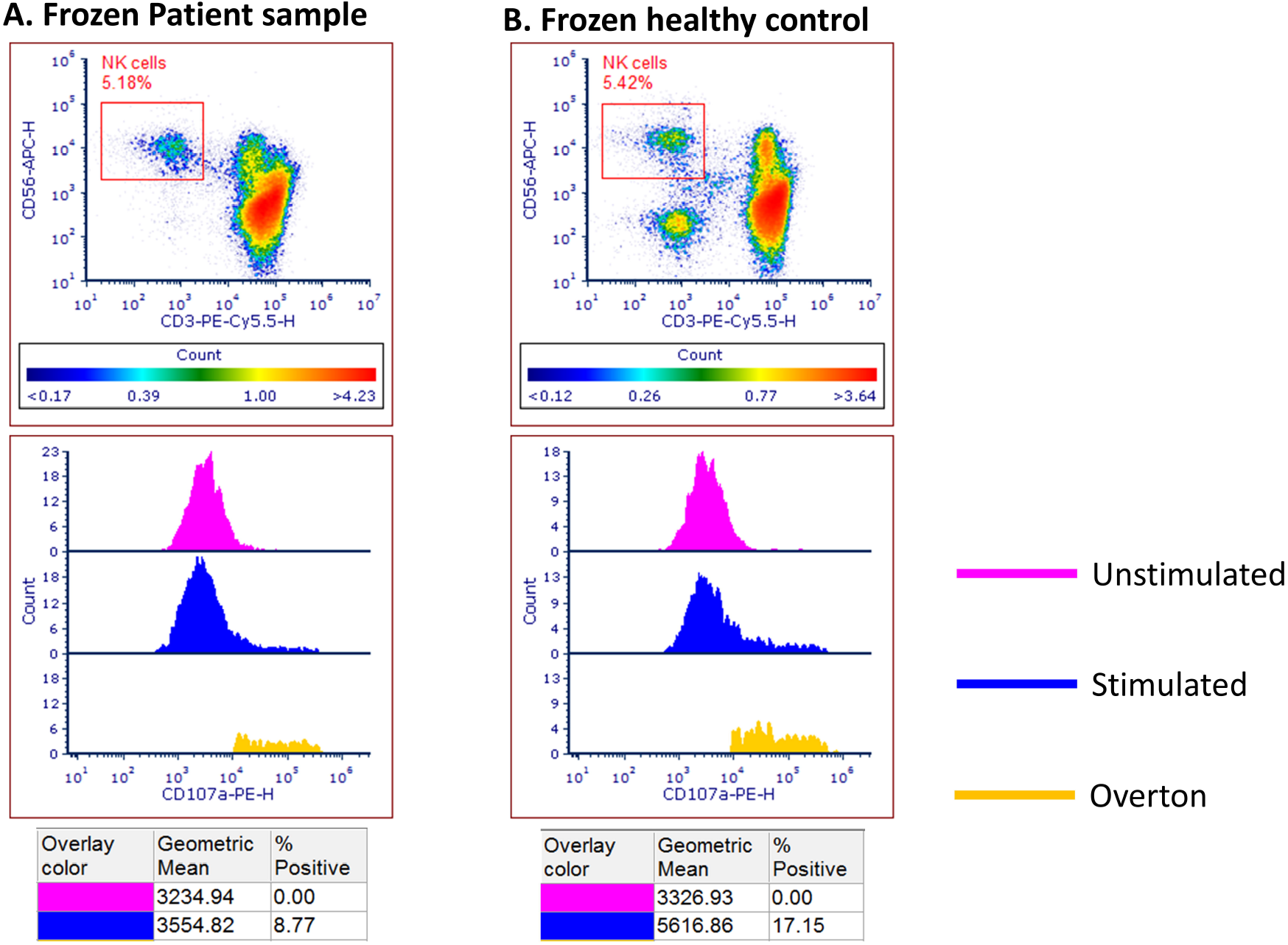
Suspected HLH case. A suspected case of HLH was evaluated for NK cell degranulation using the 18-hour time point. The 6-hour assay could not be performed due to the sample arriving over the weekend, which limited staff availability. Since the 6-hour protocol was under evaluation at CCHMC, the 18-hour incubation was selected for in-house testing. Patient PBMCs, along with a healthy control sample collected on the same day, were isolated and cryopreserved at –80°C. The following week, Day 2 testing was conducted on both frozen samples. The patient sample showed reduced NK cell degranulation (CD107a positivity = 8.77%), while both the frozen and healthy control samples exhibited CD107a expression above the 10% threshold. Rituximab treatment was associated with depletion of B cells (CD3□CD56□) in the patient.

**Figure 8.**
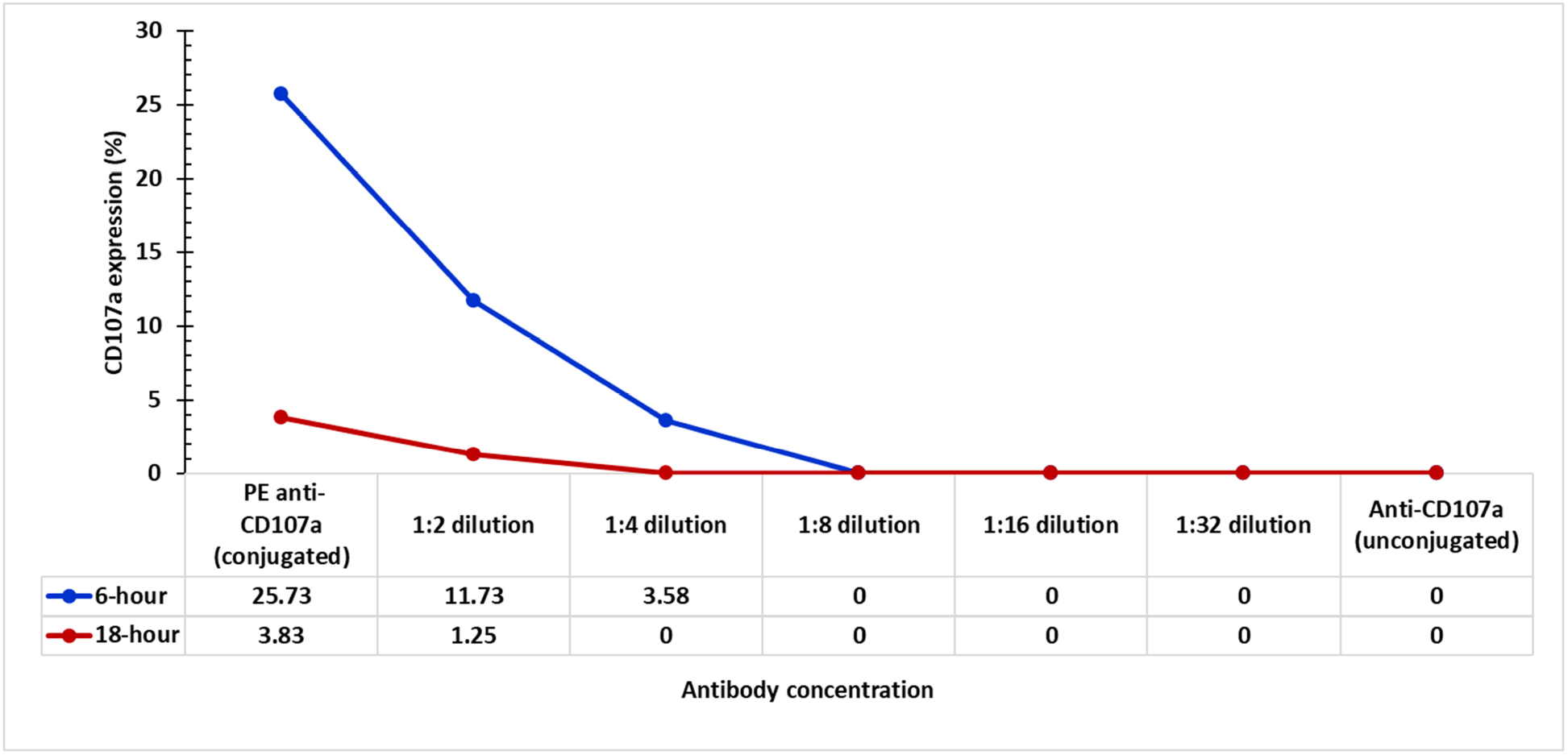
Assay sensitivity. The 6-hour protocol demonstrated a clear, dose-dependent reduction in NK cell degranulation as the ratio of unconjugated to conjugated anti-CD107a increased in the reaction mix. The highest CD107a expression (28%) was observed with 20 µL of undiluted conjugated anti-CD107a, as recommended by the manufacturer. A similar trend was noted with the 18-hour protocol; however, due to a significantly lower CD107a expression (4%) under the same undiluted condition, the resulting curve lacked distinction. Both protocols showed complete loss of CD107a positivity at a 1:8 dilution of conjugated to unconjugated anti-CD107a.

### Assay sensitivity

It was observed that manufacturer recommended volume of 20µL anti-CD107-PE (undiluted, directly from the vial) generated highest NK cell degranulation. For both 6- and 18-hour time periods. A notable decrease in the detection of surface CD107a was observed as the dilution of unconjugated anti-CD107a increased, for both 6-hour and 18-hour time periods. The decline in sensitivity was more pronounced at the 6-hour mark compared to the 18-hour mark.

## Discussion

Low or absent NK cell activity is one of the criteria for HLH diagnosis [9]. Upon exposure to foreign cells such as tumor or virus infected cells, NK cells become activated and degranulate, triggering an immune response to destroy these invading cells, in the process expressing CD107a protein on the surface [21]. Staining activated NK cells with anti-CD107a has been an established method to capture NK cell activity in HLH patients [22]. MHC-deficient K562 cells are commonly utilized as a physiological stimulus in NK cell degranulation assays [1, 3].

In this study, we tested two protocols that differed only in stimulation periods (6 hours versus 18 hours) to activate NK cells. Positive CD107a expression was determined by comparing right shift of K562 stimulated NK cell peak to unstimulated NK cell peak from the same subject. We found it challenging to draw cursors based solely on visual evaluation of histograms, which could introduce user bias. To simplify this process, we initially created density plots for the stimulated and unstimulated sets (Figure 4). To standardize data analysis and minimize human error, we automated the calculation of % positive CD107a cells using subtraction plots, which deducted unstimulated CD107a expression from stimulated CD107a expression to yield true CD107a expression on degranulated NK cells.

Our findings indicated that many samples exhibited lower CD107a expression when subjected to longer stimulation periods, as seen in 18-hour protocol. Previously published findings by Chiang et al., have demonstrated that stimulation beyond 6 hours can diminish CD107a expression on anti-CD3 stimulated cytotoxic T-lymphocytes (CTL) and anti-CD16 stimulated NK cells [16]. While our aim was to explore longer stimulation periods, the current results suggest that it is optimal to conduct NK cell degranulation assays within a 6-hour timeframe. Our findings indicate that 18-hour time frame would not be suitable for clinical diagnosis of NK cell degranulation defects. Given that K562-stimulated NK cell degranulation assays can be combined with multiple stimulants and also used to evaluate CTL degranulation, it is prudent to utilize a time point that accommodates both NK cells and CTLs.

It seems that allowing PBMCs to rest immediately after isolation aids in returning the cells to their natural, unstimulated state. However, the optimal duration for this resting period still needs further investigation. In clinical laboratories, implementing an overnight resting period has proven to be more effective, particularly in relation to staffing schedules and the workload associated with various clinical tests.

Inter-laboratory comparisons were performed only on two healthy controls and one suspected HLH case. Despite differing instrumentation and analytical protocols, no significant difference was observed in the data from both laboratories. However, the limited number of samples tested warrants further verification with a larger sample size. Although our results from frozen samples were concordant with CCHMC’s results, the assay should be evaluated with more samples to better understand the impact of sample freezing on NK cell degranulation defects.

Throughout our study, we observed moderate to high inter-day variability in CD107a expression from the same donor sample, despite no changes in the donor’s health condition (Figure 9). This variability could be attributed to the K562 cells used, which may originate from different passage numbers. Given that maintaining cell lines is not feasible for many clinical laboratories, and that a single stimulant, such as the K562 cell line, may produce variable data, some laboratories have opted to use alternative stimulants like PMA/Ionomycin and phytohemagglutinin (PHA) to assess NK cell degranulation [23–25]. While NK cells can be stimulated by K562 cells, anti-CD16, or PMA/ionomycin, T cells can be stimulated by anti-CD3 or PMA/ionomycin. In our study, we chose to focus solely on physiological stimulation to evaluate NK cell degranulation, specifically using K562 cells. Furthermore, the 6-hour assay has been traditionally optimized with K562 cells, so our initial efforts were concentrated on fine-tuning the time period specifically assessing whether an 18-hour protocol could effectively substitute the traditional 6-hour method before exploring additional stimulants.

**Figure 9.**
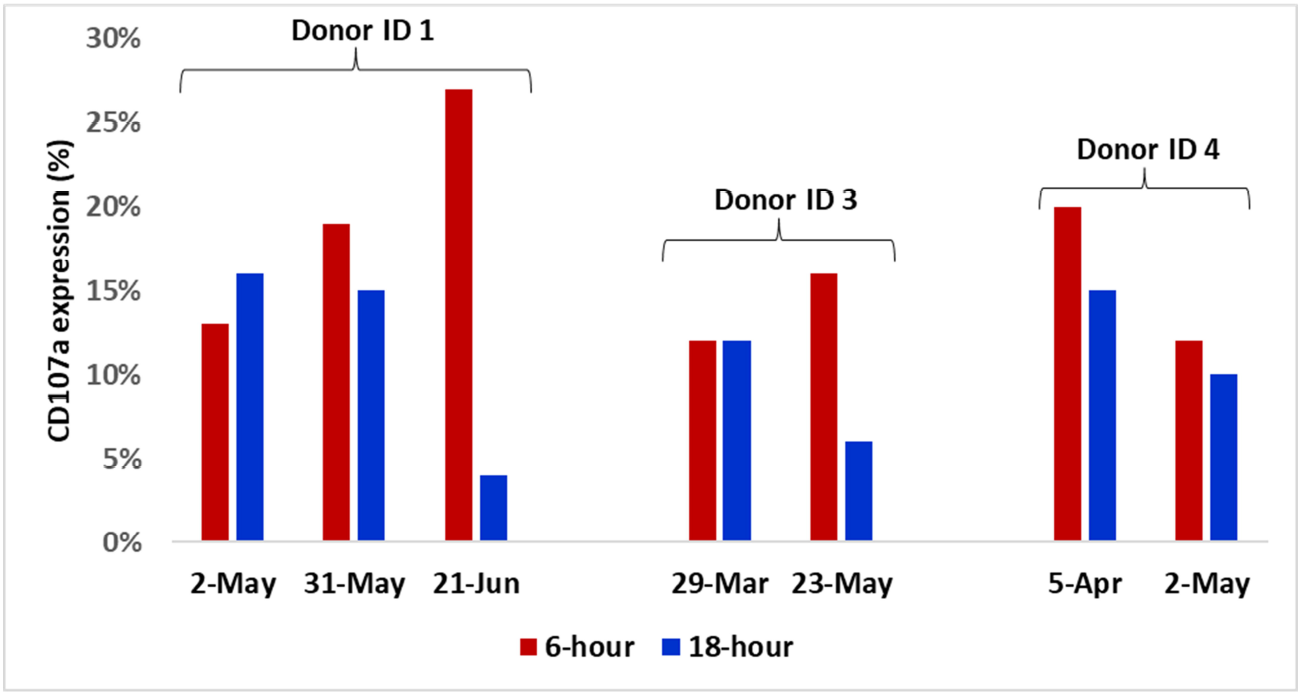
Inter-day donor variability. CD107a expression was measured in three donors using both the 6-hour and 18-hour protocols. Despite the donors maintaining stable health conditions, repeated measurements revealed variability in CD107a expression.

Combining NK cell degranulation assays with T-cell degranulation assays may improve the generation of clinical data supporting HLH diagnoses [11, 13]. For example, in familial HLH type 3, analysis of the CD3□CD8□CD57□ subset of cytotoxic T lymphocytes yielded conclusive results [8]. In another study, Vγ9Vδ2 T-cell degranulation demonstrated higher specificity than traditional NK cell degranulation assays for HLH diagnosis [26]. In developing this assay in our laboratory, we have observed that T-cell degranulation data tend to be more consistent than NK cell degranulation results, particularly when NK cells are stimulated with K562 cells. Importantly, some patients may present with low NK cell counts; in such cases, evaluating additional cell types could provide clinicians with valuable insights and support more accurate diagnoses. Thus, this multifaceted approach underscores the potential for improved clinical outcomes in HLH.

It is important to recognize that NK cell–based assay performance can be influenced by sample storage and shipping conditions. For example, we once received isolated PBMCs from a patient previously reported to have diminished NK cell degranulation. The sending laboratory had isolated PBMCs from sodium heparin whole blood within 24 hours of collection and shipped them overnight in a Styrofoam container with cold packs to maintain temperature. Upon receipt, we allowed the PBMCs to incubate overnight to facilitate recovery from potential transit stress before repeating the NK cell degranulation assay, using the same protocol as the sending laboratory. Unexpectedly, the patient’s NK cell degranulation appeared normal. We hypothesize that the shipping process itself may have inadvertently activated NK cells, producing results that differed markedly from assays performed directly on freshly isolated cells. Based on this observation, we recommend that whole blood, rather than pre-isolated PBMCs, be considered the standard specimen type for NK cell degranulation testing. To our knowledge, most U.S. centers that perform clinical NK cell degranulation assays already accept whole blood at room temperature, provided samples are processed within 24 hours of collection.

We recognize that the current study has a limited scope, a relatively small sample size, and its findings may primarily benefit clinical laboratories that routinely conduct CD107a assays or are investigating different timeframes. Additionally, because the CD107a assay is time-consuming and resource-intensive, it was not cost-effective to continue protocol comparison runs once it became clear, over a four-month period, that the 18-hour protocol did not yield data comparable to the 6-hour time point. However, it is important to note that both primary and secondary forms of HLH, though rare, are associated with a high mortality rate [27–30]. Consequently, early diagnosis of HLH is crucial in managing this debilitating illness. This underscores the necessity of advancing research methodologies aimed at improving existing testing techniques. By doing so, we can enhance our ability to diagnose and treat HLH effectively, ultimately saving lives.

## Contributions

LF performed the performed the experiments. LF, PP: analyzed the data. LF, LK, JD, MP, PP participated in critically evaluating and finalizing the data. PP wrote the manuscript. MP, PP confirmed final version of the manuscript.

## Acknowledgement

Staff, Clinical Immunology Laboratory, Children’s Hospital of Philadelphia for maintenance of K562 cell line. Casey Wells, Samuel Chiang, Diagnostic Immunology Laboratory, Cincinnati Children’s Hospital Medical Center.

## Conflict of Interest

Authors declare no conflict of interest.

## Use of AI Tools

The manuscript has original content. It was written completely by the corresponding author and not AI. To check the grammar and enhance certain portions of the manuscript, text enhancer function of tool, Originality.ai (https://originality.ai/text-enhancer) or rephrase function of tool ‘ChatGPT’ was utilized. AI rephrased sections of text were read, and language/readability was confirmed by the corresponding authors.

## Notes

### Competing Interest Statement

The authors have declared no competing interest.

